# Hyper-excitability and hyper-plasticity disrupt cerebellar signal transfer in the *IB2* KO mouse model of autism

**DOI:** 10.1101/300228

**Authors:** Teresa Soda, Lisa Mapelli, Francesca Locatelli, Laura Botta, Mitchell Goldfarb, Francesca Prestori, Egidio D’Angelo

## Abstract

Autism spectrum disorders (ASD) are pervasive neurodevelopmental conditions that often involve mutations affecting synaptic mechanisms. Recently, the involvement of cerebellum in ASD has been suggested but the underlying functional alterations remained obscure. We investigated single-neuron and microcircuit properties in IB2 KO mice, which present a cerebellar phenotype associated with ASD. Granule cells showed a larger NMDA receptor-mediated current and enhanced intrinsic excitability raising the excitatory/inhibitory balance. Furthermore, the spatial organization of granular layer responses to mossy fibers shifted from a *Mexican hat* to *stovepipe hat* profile, with stronger excitation in the core and weaker inhibition in the surround. Finally, the size and extension of long-term synaptic plasticity was remarkably increased. These results show for the first time that hyper-excitability and hyper-plasticity disrupt signal transfer in the granular layer of IB2 KO mice supporting cerebellar involvement in the pathogenesis of ASD.

## Introduction

Autism Spectrum Disorders (ASDs) are pervasive developmental disorders characterized by impairment in social communication and social interaction and by the presence of repetitive behaviors and/or restricted interests. ASDs cover a spectrum of different clinical conditions ranging from severely hypofunctional to hyperfunctional, and show abnormalities in different brain regions. Although most attention has been given so far to the cerebral cortex, increasing evidence implicates also the cerebellum (Amaral, 2011; Betancur, 2011; Ellegood et al., 2015). Cerebellar lesions often cause autistic-like symptoms (Hampson and Blatt, 2015) and perinatal cerebellar injuries are the greatest non-genetic risk factor for ASD (Bolduc and Limperopoulos, 2009; Limperopoulos et al., 2009; Bolduc et al., 2011; Wang et al., 2014; Mosconi et al., 2015). Moreover, cerebellar alterations are found in several syndromic forms of ASD, like Phelan-McDermid, Fragile X, Tuberous Sclerosis and Rett syndrome [for recent reviews, see (Courchesne and Allen, 1997; Schmahmann, 2004; Allen, 2006; Ito, 2008; D’Angelo and Casali, 2013; Broussard, 2014; Hampson and Blatt, 2015; Mosconi et al., 2015; Zeidán-Chuliá et al., 2016)]. This raises a main question: are there any alterations of cerebellar microcircuit functions in ASD?

ASDs are often associated with mutations in genes coding for synaptic proteins (Qiu et al., 2012; Banerjee et al., 2014; De Rubeis and Buxbaum, 2015; Kim et al., 2016) bringing about neurotransmission abnormalities (Curatolo et al., 2014; Ellegood et al., 2015; Kloth et al., 2015; Mercer et al., 2016; Sztainberg and Zoghbi, 2016; Tsai, 2016; Tu et al., 2017). The consequent microcircuit alterations have mainly been analyzed in the neocortex revealing that: (i) hyper-reactivity to stimulation, accompanied by altered neuronal excitability and synaptic plasticity, was related to increased glutamatergic transmission (Rinaldi et al., 2007; Markram et al., 2008; Rinaldi et al., 2008c; Markram and Markram, 2010); (ii) dysregulation of the excitatory/inhibitory (E/I) balance was related to various alterations at excitatory and inhibitory synapses (Rubenstein and Merzenich, 2003; Gogolla et al., 2009; Uzunova et al., 2015); (iii) altered modular organization of microcircuits (Casanova, 2003, 2006; Hutsler and Casanova, 2016) was related to reduced lateral inhibition, bringing about changes in the spatial organization of neuronal activation and synaptic plasticity. In particular, center-surround (C/S) structures were proposed to change from a “Mexican hat” to a “stovepipe hat” profile (Casanova, 2006).

A key role in synaptic and microcircuit dysregulation has been suggested by NMDA receptor hyperfunction and NMDA receptor antagonists have been recently reported to mitigate ASD symptoms in *Mef2c* mice models of Rett syndrome (Tu et al., 2017). Important for the present case, NMDA receptor-mediated currents were increased in cerebellar granule cells of the IB2 (Islet Brain-2) KO mouse, a model of the Phelan-McDermid syndrome (Giza et al., 2010). IB2 (MAPK8IP2) is a scaffolding protein enriched in the PSD, probably regulating signal transduction by protein kinase cascades, that operates inside the NMDA receptor interactome (Yasuda et al., 1999). Since NMDA receptor expression in granule cells is the highest among cerebellar neurons (Monaghan and Cotman, 1985) and has a profound impact on synaptic excitation and plasticity (D’Angelo et al., 1995; Armano et al., 2000; Sola et al., 2004; D’Errico et al., 2009), IB2 KO mice actually provide an ideal model to investigate cerebellar microcircuit alterations in ASD. In the cerebellar granular layer, granule cells receive excitatory synapses from mossy fibers and are inhibited by Golgi cells. The synaptic interaction between these neurons forms the granular layer microcircuit which, once activated by incoming spike bursts, generates responses organized in C/S (Mapelli and D’Angelo, 2007; Gandolfi et al., 2014). Here we show that the granular layer of IB2 KO mice is characterized by hyper-excitability and hyper-plasticity, which raise the E/I balance disrupting C/S structures and signal transfer at the input stage of cerebellum. The implications of these cerebellar microcircuit alterations for ASD pathogenesis are discussed.

## Methods

All procedures were conducted in accordance with European guidelines for the care and use of laboratory animals (Council Directive 2010/63/EU) and approved by the Ethical Committee of Italian Ministry of Health (637/2017-PR).

### Genotyping and maintenance of IB2 KO mice

Experiments were conducted on IB2^+/+^ (WT) and IB2^-/-^ (KO) mice obtained by crossing IB2^+/-^ parents, since IB2 KO are poor breeders, possibly reflecting the social deficit associated with IB2 deletion (Giza et al., 2010). The genotyping was conducted through PCR using four primers to detect wild-type and null alleles as previously described (Giza et al., 2010).

### Slice preparation and solutions

The experiments reported in this paper have been conducted on 17-to 24-day-old (P0=day of birth) WT and IB2 KO mice. Mice were anesthetized with halothane (Sigma, St.Louis, MO) and killed by decapitation in order to remove the cerebellum for acute slice preparation according to a well-established technique (D’Angelo et al., 1995; Armano et al., 2000; Gall et al., 2005; Prestori et al., 2013; Nieus et al., 2014). The vermis was isolated and fixed on the vibroslicer’s stage (Leica VT1200S) with cyano-acrylic glue. Acute 220 μm-thick slices were cut in the parasagittal plane in ice cold (2–3°C) Krebs solution containing (in mM): 120 NaCl, 2 KCl, 1.2 MgSO_4_, 26 NaHCO_3_, 1.2 KH_2_PO_4_, 2 CaCl_2_, and 11 glucose, equilibrated with 95% O_2_-5% CO_2_ (pH 7.4). Slices were allowed to recover at room temperature for at least 1h, before being transferred to a recording chamber mounted on the stage of an upright microscope (Zeiss, Oberkochen, Germany). The slices were perfused with oxygenated Kreb’s solution and maintained at 32°C with a Peltier feedback device (TC-324B, Warner Instrument Corp., Hamden, CT). For electrophysiological recordings, Kreb’s solution was added with the GABA_A_ receptor antagonist SR95531 (gabazine, 10 μM; Sigma). In some experiments, Kreb’s solution was Mg^2+^ -free. Local perfusion with Krebs solution and 10 μM SR95531 was commenced before seal formation and was maintained until end of recording. In a set of experiments the GABA_A_ receptor antagonist SR95531 was omitted from the Krebs solution.

### Electrophysiological recordings

Whole-cell patch-clamp recordings were performed with Multiclamp 700B [-3dB; cutoff frequency (fc),10 kHz], sampled with Digidata 1440A interface, and analyzed off-line with pClamp10 software (Molecular Devices, CA, USA). Patch pipettes were pulled from borosilicate glass capillaries (Sutter Instruments, Novato, CA) and filled with different solutions depending on the specific experiments (see below). Mossy fiber stimulation was performed with a bipolar tungsten electrode (Clark Instruments, Pangbourne, UK) via a stimulus isolation unit. The stimulating electrode was placed over the central fiber bundle in the cerebellar lamina to stimulate the mossy fibers, and 200 μs step current pulses were applied at the frequency of 0.1-0.33 Hz (in specific experiments, paired-pulse stimulation at 20 ms inter-pulse was used). From a comparison with data reported in (Sharma and Vijayaraghavan, 2003; Giza et al., 2010; Sgritta et al., 2017), 1 or 2 mossy fibers were stimulated per granule cell in the experiments used for quantal analysis. Long-term potentiation (LTP) induction was obtained by a continuous stimulation of 100 pulses at 100Hz at –50 mV (HFS), as reported previously (Armano et al., 2000; Gall et al., 2005; D’Errico et al., 2009; Prestori et al., 2013). Results are reported as mean ± SEM and compared for their statistical significance by unpaired Student’s test (unless otherwise stated; a difference was considered significant at p < 0.05).

The stability of whole-cell recordings can be influenced by modification of series resistance (R_s_). To ensure that R_s_ remained stable during recordings, passive electrode-cell parameters were monitored throughout the experiments. The granule cell behaves like a lumped electrotonic compartment and can therefore be treated as a simple resistive - capacitive system, from which relevant parameters can be extracted by analyzing passive current relaxation induced by step voltage changes. In each recording, once in the whole-cell configuration, the current transients elicited by 10 mV hyperpolarizing pulses from the holding potential of −70 mV in voltage-clamp mode showed a biexponential relaxation, with a major component related to a somatodendritic charging (Prestori et al., 2008). According to previous reports (D’Angelo et al., 1995; Silver et al., 1996; D’Angelo et al., 1999), the major component was analyzed to extract basic parameters useful to evaluate the recordings conditions and to compare different cell groups. Membrane capacitance (C_m_) was measured from the capacitive charge (the area underlying current transients) and series resistance was calculated as R_s_ = τ_vc_/C_m_. The membrane resistance (R_m_) was computed from the steady-state current flowing after termination of the transient. The 3-dB cut-off frequency of the electrode-cell system was calculated as f_vc_ = (2π • τvc)^-1^. The data are reported in Table 1. In the cells considered for analysis, these values did not significantly change after 30 minutes attesting recording stability. Cells showing variation of series resistance (R_s_) >20% were discarded from analysis.

**Table 1.** Properties of whole-cell recordings in mice granule cells. The data were obtained using K-gluconate intracellular solution and analyzing current transient elicited by 10 mV voltage-clamp steps delivered from the holding potential of −7 0mV. The number of observations indicated and statistical significance is reported in comparison with IB2 KO granule cells. ^∗∗^p<0.01, unpaired t test.

|  | WT<br>(n = 70) | IB2 KO<br>(n = 57) |
| --- | --- | --- |
| $R_m$ (GΩ) | 2.3 ± 0.1 | 2.9 ± 0.3 |
| $C_m$ (pF) | 3.6 ± 0.09 | 3.2 ± 0.09 ** |
| $R_s$ (MΩ) | 15.9 ± 0.9 | 16.9 ± 1.0 |
| $f_{vc}$ (KHz) | 3.8 ± 0.3 | 3.5 ± 0.2 |
| $\tau_{vc}$ (μs) | 54.1 ± 3.0 | 60.5 ± 7.2 |
| $V_m$ (mV) | -52.1 ± 2.6 (n=12) | -54.5 ± 2.0 (n=12) |

### Granule cell excitability

Patch pipettes had 7-9 MΩ resistance before seal formation with a filling solution containing (in mM): 126 potassium gluconate, 4 NaCl, 5 Hepes, 15 glucose, 1 MgSO_4_.7H_2_O, 0.1 BAPTA-free, 0.05 BAPTA-Ca^2+^, 3 Mg^2+^-ATP, 0.1 Na -GTP, pH 7.2 adjusted with KOH. The calcium buffer is estimated to maintain free calcium concentration around 100 nM. Just after obtaining the cell-attached configuration, electrode capacitance was carefully cancelled to allow for electronic compensation of pipette charging during subsequent current-clamp recordings. At the beginning of each recording, a series of depolarizing steps was applied in voltage-clamp to measure the total voltage-dependent current of the granule cell (see Fig. 1C). Leakage and capacitance were subtracted using a hyperpolarizing pulses delivered before the test pulse (P/4 protocol). After switching to current-clamp, intrinsic excitability was investigated (see Fig. 1B) by setting resting membrane potential at −80 mV and injecting 800-ms current steps (from - 4 to 22 pA in 2 pA increment). Membrane potential during current steps was estimated as the average value between 600 and 800 ms. Action potential frequency was measured by dividing the number of spikes by step duration.

**Figure 1.**
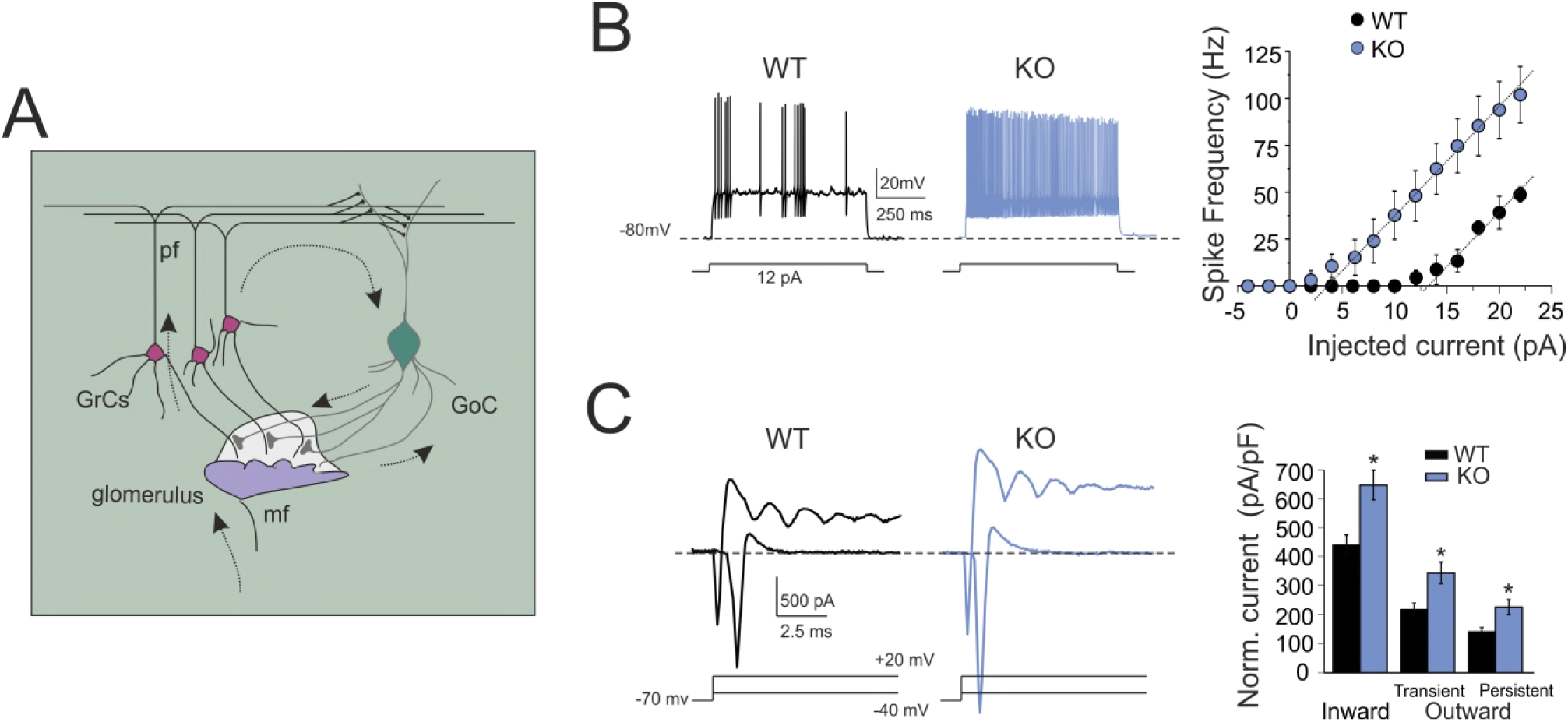
Granule cell excitable properties. A. Schematic representation of cerebellar circuit. Mossy fibers (mf) contact granule cells (GrC) and Golgi cell (GoC) dendrites. Axons of GrCs, the parallel fibers (pf), activate Golgi cells which inhibit GrCs through feedforward and feedback inhibitory loops.
B. Granule cell electroresponsiveness. Voltage responses were elicited from −80 mV using step current injection. The plot shows the relationships between average spike frequency over 2 sec and the injected current intensity both for WT (n=6) and IB2 KO (n=8) granule cells. Linear fits (dashed lines): WT x-intercept 3 pA, slope 7.8 ± 1.6 spike/pA (n=6); IB2 KO x-intercept 13 pA, slope 6.4 ± 0.5 spike/pA (n=8). Data are reported as mean ± SEM.
C. Voltage-activated inward and outward currents in granule cells. Exemplar voltage-dependent currents evoked by depolarizing voltage steps from the holding potential of −70 mV were leak-subtracted. The histogram compares inward and outward current amplitudes measured at −40 mV and +20 mV in WT and IB2 KO mice. Data are reported as mean ± SEM; ^∗^p<0.05.

### Post-synaptic currents

Patch pipettes had 5–8 MΩ resistance before seal formation with a filling solution containing the following (in mM): 81 Cs_2_SO_4_, 4 NaCl, 2 MgSO_4_, 1 QX-314 (lidocaine N-ethyl bromide), 0.1 BAPTA-free and 0.05 BAPTA-Ca^2+^, 15 glucose, 3 Mg^2+^-ATP, 0.1 Na^+^-GTP, and 15 HEPES, pH adjusted to 7.2 with CsOH. The calcium buffer is estimated to maintain free calcium concentration around 100 nM. Synaptic currents elicited at 0.33 Hz were averaged and digitally filtered at 1.5 kHz off-line. IPSC and EPSC peak amplitude were taken at +10 and −70 mV to measure the GABA_A_ and AMPA currents, respectively. In some experiments, NMDA current was directly measured at - 70 mV in Mg^2+^ -free solution in the presence of the AMPA receptor blocker, 10 μM NBQX (Sola et al., 2004). In LTP experiments, the acquisition program automatically alternated EPSC with background activity recordings (1 s and 9 s, respectively), from which mEPSCs were detected. After 10 min (control period), the recording was switched to current clamp (patch pipettes were filled with a K^+^-gluconate based solution) and high-frequency stimulation (HFS) was delivered to induce plasticity. Long-term synaptic efficacy changes were measured after 20 min. After delivering HFS, voltage-clamp at −70 mV was reestablished and stimulation was restarted at the test frequency. EPSCs and mEPSCs were digitally filtered at 1.5 kHz and analyzed off-line with pClamp10 software (Molecular Devices, Sunnyvale, CA). For both EPSC and mEPSC peak amplitude was computed. mEPSC detection was performed automatically with Mini Analysis Program (Synaptosoft, Inc. Decatur, GA) when their amplitude was 5-7 time the baseline noise S.D. (0.88 ± 0.03; n=8). These criteria and a further visual inspection of detected signals allowed us to reject noise artifacts.

In order to investigate the expression mechanism of long-term synaptic plasticity over a heterogeneous data set (Sola et al., 2004; Gall et al., 2005), a simplified version of quantal analysis was performed by measuring the mean (M) and standard deviation (S) of EPSC amplitude. EPSC changes, which do not strictly require that single synaptic connections are isolated, were obtained from M and S: the coefficient of variation, CV = S/M, the paired-pulse ratio, PPR = M_2_/M_1_, i.e. the ratio between the second and first EPSC amplitude in a doublet at 20 ms inter-pulse interval. The comparison between M and CV obtained before and after the induction of plasticity could be performed in the plot (CV_2_/CVA_1_)^-2^ *vs*. (M_2_/M_1_). Assuming binomial statistics, this plot has the property that the unitary slope diagonal separates points caused by changes in quantum content (m = *np*, with *n* being the number of releasing sites and *p* the release probability) from those caused by changes in quantum size (*q*). The inequality leads to a topological representation of neurotransmission changes (see Fig.7) and has been extensively used to interpret the plasticity mechanism (Bekkers and Stevens, 1990; Malinow and Tsien, 1990; Sola et al., 2004; Rinaldi et al., 2008a; D’Errico et al., 2009; Sgritta et al., 2017). For an M increase:

i. when (CV_2_/CVA_1_)^−2^ > (M_2_/M_1_) both *n* and *p* can increase,
ii. when (CV_2_/CVA_1_)^−2^ = (M_2_/M_1_) only *n* can increase,
iii. when (CV_2_/CVA_1_)^−2^ < (M_2_/M_1_) neither *n* nor *p* can increase implying an increase in *q.* A pure increase in *q* will lie on the axis when (CV_2_/CVA_1_)^-2^ =1.

### Voltage sensitive dye imaging (VSDi)

The stock solution for VSDi contained the dye Di-4-ANEPPS (Molecular Probes, Eugene, OR) dissolved in a Krebs-based solution containing 50% ethanol (Sigma) and 5% Cremophor EL (Sigma). Slices for optical recordings were incubated for 30 minutes in oxygenated Krebs solution added with 3% Di-4-ANEPPS stock solution and mixed with an equal volume of fetal bovine serum (Molecular Probes) to reach a final dye concentration of 2 mM (Vranesic et al., 1994). After incubation, the slices were rinsed with Krebs solution to wash out the dye that was not incorporated by the tissue, before being transferred to the recording chamber installed on an upright epifluorescence microscope (Slicescope, Scientifica Ltd, Uckfield, UK), equipped with a 20X objective (XLUMPlanFl 0.95 NA, water immersion; Olympus, Tokyo, Japan). The light generated by a halogen lamp (10V150W LM150, Moritex, Tokyo, Japan) was controlled by an electronic shutter (Newport corporation, Irvine, CA) and then passed through an excitation filter (λ = 535 ± 20 nm), projected onto a dichroic mirror (λ = 565 nm) and reflected toward the objective lens to illuminate the specimen. Fluorescence generated by the tissue was transmitted through an absorption filter (λ > 580 nm) to the CCD camera (MICAM01, Scimedia, Brainvision, Tokyo, Japan). The whole imaging system was connected through an I/O interface (Brainvision) to a PC controlling illumination, stimulation and data acquisition. The final pixel size was 4.5×4.5μm with 20X objective. Full-frame image acquisition was performed at 0.5 kHz. Data were acquired and displayed by Brainvision software and signals were analyzed using custom-made routines written in MATLAB (Mathworks, Natick, MA). At the beginning of recordings, a calibration procedure was adopted to ensure homogeneity across experiments. The dynamic range of the CCD camera was calibrated by measuring background fluorescence and setting the average light intensity in the absence of stimulation to 50% of the saturation level. The background fluorescence was sampled for 50 ms before triggering electrical stimulation and was used to measure the initial fluorescence intensity (*F*_0_). The relative fluorescence change (Δ*F*/*F*_0_) was then calculated for each time frame. The signal-to-noise ratio was improved by averaging 10 consecutive sweeps at the stimulus repetition frequency of 0.1 Hz.

### VSDi data analysis

Fluorescence data collected by Brainvision acquisition software were filtered using both a cubic filter (3×3) and a spatial filter (3×3) embedded in the software, and then exported and processed in Matlab. The resulting files were a series of matrices each representing a temporal frame of the acquired trace. Using appropriate Matlab routines written *ad hoc*, single matrices representing the peak value of granular layer responses to electrical stimulation were obtained. These maps containing the information on the signal peak amplitudes and their spatial origin were used for comparison of control condition and different treatments, as detailed below. Data were reported as mean ± SEM. Statistical significance was assessed using unpaired Student’s t test unless otherwise stated. For the analysis of the amount and spatial distribution of the NMDA receptor component of excitation in the cerebellar granular layer of WT and IB2 KO mice, responses to electrical stimulation of the mossy fibers were recorded in control and after perfusion of the NMDA receptor blocker APV (50 μM). The average map of APV effect on signal amplitudes was subtracted to the control map, to unveil the contribution of the NMDA receptors. The spatial distribution of the NMDA receptor-mediated depolarization was revealed by averaging each experimental map on the peak of NMDA receptor component in each case. Whenever spatial maps obtained from different experiments were averaged, the corresponding slices were aligned along the mossy fiber bundle. For the analysis of the excitatory/inhibitory (E/I) balance and spatial distribution of excitation and inhibition in the granular layer, similar experiments were carried out, recording the responses to MFs stimulation before and after the perfusion of the GABA_A_ receptor antagonist SR95531 (gabazine; 10 μM). This approach allows to reconstruct a map of regions with prevailing excitation (E) compared to regions showing prevailing inhibition (I) (Mapelli and D’Angelo, 2007; Gandolfi et al., 2014). In this case, the E map was constructed on the control responses (where the response is available only in the regions where excitation prevails over inhibition), while the I map was constructed subtracting the maps after SR95531 perfusion to the control maps (unveiling the regions where, before SR95531 perfusion, excitation was prevented by inhibition). Both E and I maps were normalized to 1, and the E/I balance maps were obtained as (E-I)/E. The C/S organization of excitation and inhibition was evident averaging the E/I maps in each experiment on the peak of excitation in controls. For the analysis of the amount and spatial distribution of LTP and LTD in the granular layer, plasticity maps were obtained by comparing responses amplitudes before and after the plasticity induction through a HFS delivered to the mossy fiber bundle. The C/S spatial organization of LTP and LTD was unveiled by averaging each plasticity maps from different experiments on the peak of maximum LTP.

## Results

In the cerebellum granular layer, there are three main mechanisms controlling the E/I balance of granule cells (Nieus et al., 2014): granule cells intrinsic excitability, mossy fiber glutamatergic excitation, Golgi cell GABAergic inhibition (Mapelli et al., 2014) (Fig.1A). Here, these properties have been compared in turn between IB2 KO and WT mice. In patch-clamp whole-cell recordings in acute cerebellar slices, there were no significant differences in either series resistance (R_s_), membrane resistance (R_m_), or resting membrane potential between IB2 KO and WT cerebellar granule cells (Table 1).

### Enhanced intrinsic excitability in IB2 KO granule cells

In whole-cell current-clamp recordings, both WT and IB2 KO granule cells were silent at rest and responded to current steps with fast repetitive spike discharges that increased their frequency almost linearly with stimulus intensity (D’Angelo et al., 1995; Brickley et al., 1996; D’Angelo et al., 1998; Rossi et al., 1998; Armano et al., 2000; Cathala et al., 2003; Prestori et al., 2008) (Fig. 1B). However, IB2 KO granule cells showed higher discharge frequency compared to WT granule cells both at low current injection [12 pA: WT = 4.1 ± 0.1 Hz (n=6); IB2 KO = 48.1 ± 14.2 Hz (n=8); p=0.017, unpaired *t*-test] and at high current injection [20 pA: WT = 39.2 ± 9.5 Hz (n=6); IB2 KO = 93.7 ± 16.0 Hz (n=8); p=0.014, unpaired *t*-test], shifting the frequency-intensity plot toward the left (Fig. 1B). It should be noted, as explained above and in Table 1, that the enhanced intrinsic excitability in IB2 KO mice did not depend either on passive or resting properties, which did not significantly differ in the two cell groups used in these experiments.

In the same experiments, whole-cell currents elicited by depolarizing voltage steps differed in WT and IB2 KO granule cells (Fig. 1C). The “transient inward current” (corresponding to a fast Na^+^ current) (Magistretti et al., 2006) was significantly larger in IB2 KO compared to WT granule cells. The “transient and persistent outward currents” (comprising A-type, delayed rectifier, and calcium-dependent K^+^ currents) (Bardoni and Belluzzi, 1994) were also significantly larger in IB2 KO compared to WT granule cells. Thus, the enhancement of intrinsic excitability in IB2 KO granule cells was correlated with abnormal expression of voltage-dependent membrane currents.

### Similar AMPA and GABA_A_ but increased NMDA receptor mediated currents at IB2 KO granule cell synapses

Mossy fiber stimulation is known to elicit EPSCs directly through mossy fiber activation and IPSCs indirectly through activation of Golgi cells (cfr. Fig. 1A) (Cathala et al., 2003; Cesana et al., 2013; Nieus et al., 2014). Postsynaptic currents were recorded from granule cells both at −70 mV and +10 mV in order to isolate the excitatory (EPSC) from inhibitory (IPSC) component. This technique was reported previously (Mapelli et al., 2009; Nieus et al., 2014). It should be noted that, at −70 mV, NMDA receptor-mediated currents are blocked by Mg^2+^, so that the EPSC is almost purely AMPA receptor-mediated. In the present experiments, the AMPA-EPSC peak (WT = −38.1 ± 7.1 pA, n=13 vs. IB2 KO = −34.1 ± 5.7, n=7; p=0.66) and the GABA_A_-IPSC peak (WT = 45.4 ± 8.4 pA, n=13 vs. IB2 KO = 51.7 ± 13.4, n=7; p=0.69) showed similar amplitude in WT and IB2 KO mice (Fig. 2A). Accordingly, no differences were observed in the AMPA-EPSC/GABA_A_-IPSC ratio in granule cells (WT = 0.95 ± 0.15, n=13 vs. IB2 KO = 0.86 ± 0.19, n=7; p=0.71; Fig. 2B).

**Figure 2.**
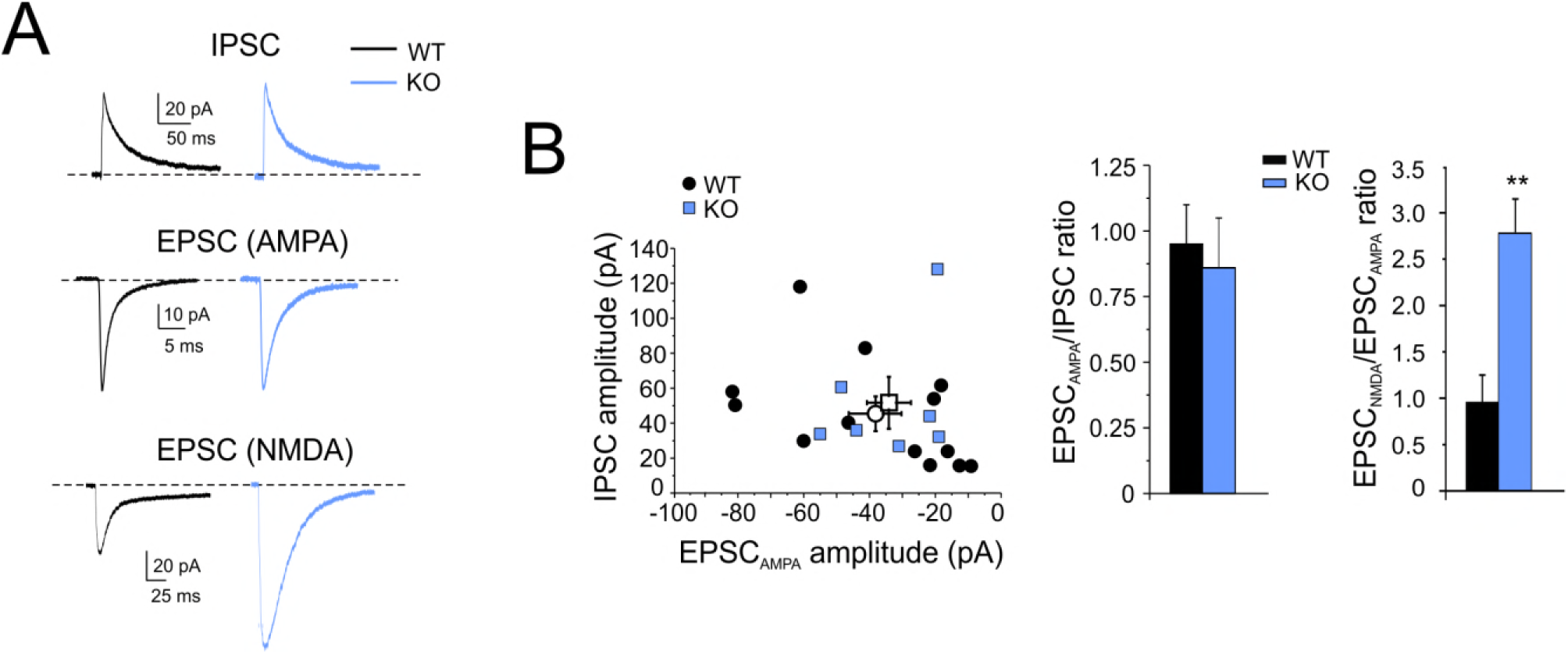
Evoked excitatory and inhibitory currents in granule cells. A. Synaptic currents in WT and IB2 KO granule cells. The EPSC_AMPA_ and IPSC are recorded from the same cells at the holding potential of −70 mV (averaging of 100 consecutive traces) and at + 10 mV (averaging 10 consecutive traces), respectively. The EPSC_NMDA_ are recorded in different cells at −70 mV in Mg^2+^-free extracellular solution in presence of the AMPA receptor antagonist, 10 μM NBQX (averaging of 30 consecutive traces).
B. IPSC/EPSC ratios at mossy fiber–granule cell synapses in WT and IB2 KO mice. The plot shows the amplitude of EPSC_AMPA_ and IPSC in the same cells for WT and IB2 KO mice (open symbols are mean ± SEM). The histogram compares the average EPSC_ampa_/IPSC ratio and EPSC_nmda_/IPSC ratio in WT and IB2 KO mice (mean ± SEM; ^∗∗^p<0.01).

In a different series of recordings, the NMDA EPSC was elicited in isolation at −70 mV in Mg^2+^ -free solution in the presence of AMPA and GABA_A_ receptor blockers (10 μM NBQX and 10 μM SR95531, respectively; Fig. 2A). The NMDA-EPSC peak was enhanced in IB2 KO synapses (WT = −37.0 ± 5.1 pA, n=6 vs. IB2 KO = −95.3 ± 17.7, n=5; p=0.03) by 2.5 times. These results confirm the alteration in NMDA EPSC amplitude reported previously (Giza et al., 2010).

In aggregate, the similarity of the AMPA-EPSC and GABA_A_-IPSC, along with the large increase of the NMDA-EPSC, suggest that the excitatory/inhibitory (E/I) balance in IB2 KO mice will move in favor of excitation in conditions in which the NMDA channels are physiologically unblocked by depolarization.

### Increased excitation in C/S structures of IB2 KO granular layer are driven by NMDA currents

In order to obtain a physiological assessment of the E/I balance and of the NMDA current contribution, we used voltage-sensitive dye imaging (VSDi). VSDi allows to generate maps of electrical activity and to investigate the spatial distribution of granular layer responses following mossy fiber stimulation (Mapelli et al., 2010). In particular, VSDi, coupled with selective pharmacological blockade of synaptic receptors, can reveal the relative role of synaptic inhibition and of NMDA receptors (Gandolfi et al., 2015).

A first set of VSDi recordings was performed by subtracting control activity maps from those obtained after GABA_A_ receptor blockade with 10 μM SR95531 (Fig. 3; see Methods for details). In agreement with previous observations, the granular layer response to mossy fiber stimuli self-organized in center/surround (C/S) structures characterized by a “Mexican hat” profile, with an excitation core surrounded by inhibition (Mapelli and D’Angelo, 2007; Solinas et al., 2010; Gandolfi et al., 2014; Gandolfi et al., 2015) (Figs 3A,B). The C/S distribution was maintained in the IB2 KO granular layer but with striking differences. (i) Excitation was enhanced generating larger cores compared to WT (core diameter: WT = 12.9 ± 1.7 μm vs. IB2 KO = 29.5 ± 4.9 μm, n=5 for both; p=0.0106) (Fig. 3C). (ii) Inhibition was weaker in the surround (WT/KO ratio I_WT/KO_ = 2.83 ± 0.17, n=5). (iii) Granular layer areas showing excitation were consequently larger in IB2 KO than WT mice (WT = 49.9 ± 3.1% vs. IB2 KO = 58.8 ± 2.1%, n=5 for both; p=0.0468; Fig. 3C). As a result, the altered C/S organization in IB2 KO showed larger excitation cores with poor inhibitory surrounds, shifting from “Mexican hat” to the so-called “stovepipe hat” shape (see Fig. 3B).

**Figure 3.**
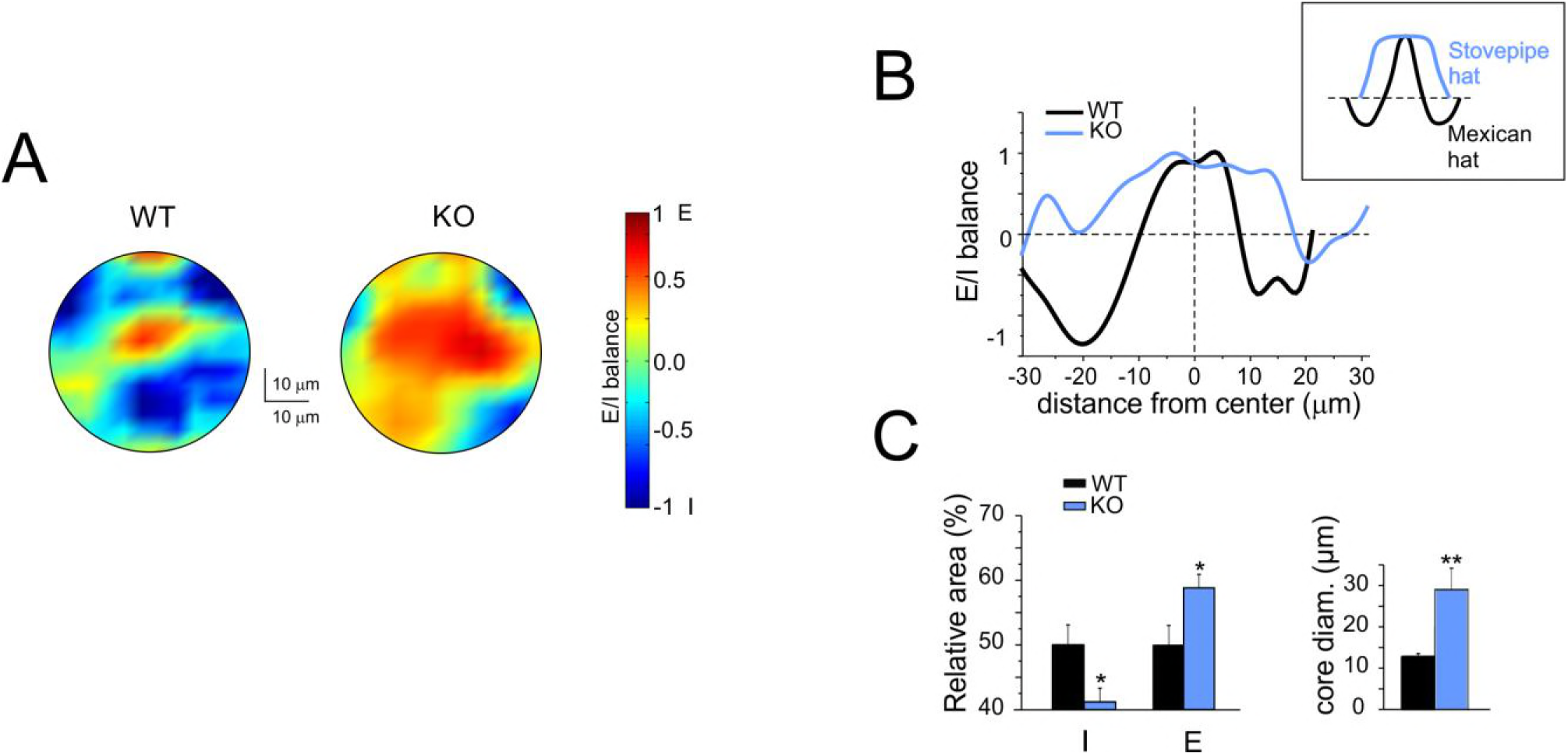
Excitatory/inhibitory balance and center/surround organization in the granular layer. A. VSDi normalized maps showing the spatial distribution of excitation and inhibition in WT and IB2 KO granular layer (average of 5 recordings in both cases).
B. The plot shows the E/I balance as a function of distance from the center for the maps shown in A. Note that in IB2 KO granular layer the excitation core is broader, while the inhibited surround is reduced, compared to WT. This tends to change the C/S from the typical Mexican hat in control to stovepipe hat shape in IB2 KO mice (cf. inset).
C. The histograms show, in WT and IB2 KO mice, the average values of the inhibition or excitation areas and of core diameter (mean ± SEM; ^∗^p<0.05, ^∗∗^p<0.01).

A second set of VSDi recordings was performed by subtracting control activity maps from those obtained after NMDA receptor blockade with 50 μM APV (Fig. 4; see Methods for details). As expected from the increased NMDA receptor-mediated current reported in Fig. 2, the NMDA receptor-mediated component of the VSDi signal was larger in IB2 KO than WT granular layers (ratio KO/WT = 2.16 ± 0.29, n=5 for both). The maps showing the spatial organization of the NMDA receptor contribution to the excitatory response were similar to the C/S organization shown in Fig. 3, with peaks of NMDA receptor contribution in cores with a diameter of 26.1 ± 1.7 μm in IB2 KO vs. 18.9 ± 1.6 μm in WT; n=5 for both; p=0.015 (Figs. 4A,B). Interestingly, since during VSDi membrane potential remains unclamped allowing voltage-dependent NMDA channel unblock during depolarization, these maps provide information about the non-linear contribution of NMDA currents. This result supported the hypothesis that the enhanced NMDA receptor-mediated transmission revealed in Fig. 2 was indeed a key player in determining the C/S and E/I alteration in IB2 KO granular layer.

**Figure 4.**
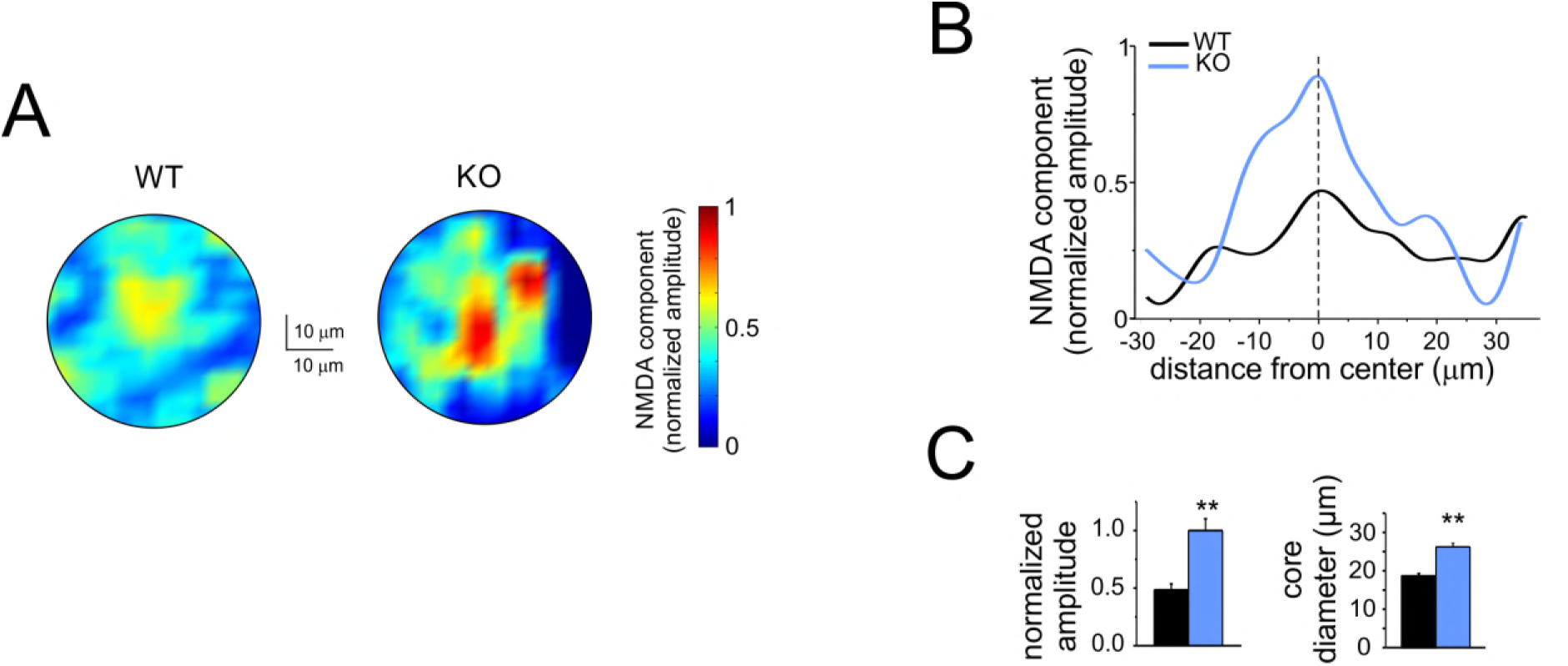
NMDA receptor-dependent component of granular layer excitation. A. VSDi normalized maps showing the spatial distribution of the NMDA component of excitation in WT and IB2 KO granular layer (average of 5 recordings in both cases).
B. The plot shows the NMDA component as a function of distance from the center for the maps shown in A. Note that in IB2 KO granular layer the NMDA component of excitation is larger and more extended compared to WT.
C. The histograms show, in WT and IB2 KO mice, the average values of the NMDA component normalized amplitude and of core diameter (mean ± SEM; ^∗∗^p<0.01).

### Enhanced long-term potentiation at the IB2 KO mossy fiber-granule cell relay

Mossy fiber-granule cell LTP is NMDA receptor-dependent through the synaptic control of postsynaptic intracellular calcium elevation (D’Angelo et al., 1999; Maffei et al., 2003; Gall et al., 2005; D’Errico et al., 2009). The impact of elevated NMDA receptor-dependent neurotransmission on LTP induction in IB2 KO mice was evaluated using a continuous high-frequency stimulation train (HFS; Fig. 5A) delivered from the holding potential of −50 mV in current-clamp (Gall et al., 2005; D’Errico et al., 2009). During HFS, IB2 KO generated more spikes than WT granule cells (WT = 23.5 ± 5.3, n=12 vs. IB2 KO = 54.2 ± 11.4, n=9; p=0.015; Figs. 5A,B), in line with the enhancement in NMDA currents (D’Angelo et al., 2005) and in intrinsic firing reported above (cf. Figs 1 and 2). After HFS, the changes were evaluated over at least 25 min after HFS.

**Figure 5.**
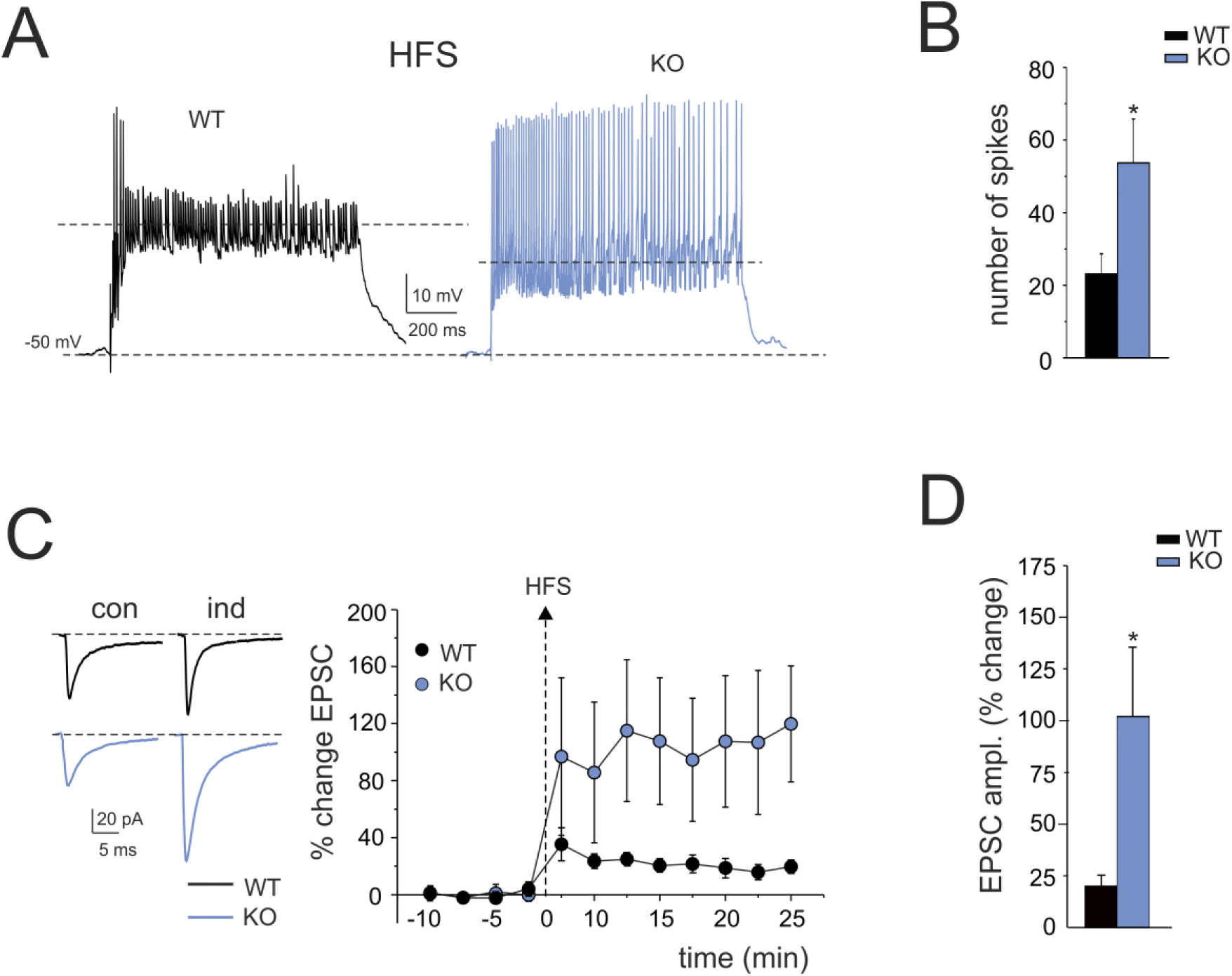
LTP of mossy fiber-granule cell EPSCs. A. Granule cell synaptic responsiveness. Voltage responses were elicited from −50 mV during 1sec-100Hz synaptic stimulation (HFS) used for plasticity induction. Note stronger spike generation in IB2 KO than WT.
B. The histogram shows the average number of spikes during HFS in WT and IB2 KO mice. Data are reported as mean ± SEM; ^∗^p<0.05.
C. LTP of EPSC_AMPA_ in WT and IB2 KO granule cells (average of 100 tracings in both cases) recorded in control and 20 min after HFS. Note that, after HFS stimulation, the EPSC_AMPA_ increase was larger in IB2 KO than WT. The LTP plot shows the average time course of EPSC_AMPA_ amplitude changes in WT (n=12) and IB2 KO (n=9) granule cells. Data are reported as mean ± SEM; ^∗^p<0.05).
D. The histogram shows the average EPSC_AMPA_ LTP following HFS in WT and IB2 KO mice. Data are reported as mean ± SEM; ^∗^p<0.05.

The AMPA EPSC increased both in WT and IB2 KO mice and remained potentiated throughout the recordings (Fig. 5C). The increase in amplitude of AMPA-EPSCs was ~5-fold larger in IB2 KO than WT mice (WT = 20.4 ± 4.2 %, n=12 vs. IB2 KO = 102.4 ± 34.9 %, n=9; p=0.047; Fig. 5D).

Intrinsic excitability increased more in WT than in IB2 KO mice (Fig. 6A,B). The current needed to generate spikes (current threshold) decreased significantly compared to control in WT granule cells (-42.8 ± 7.7%, n=6; p=0.0055) but not in IB2 KO granule cells (-8.6 ± 14.4%, n=8; p=0.07; Fig. 6B). Moreover, the increase in spike frequency was less pronounced in IB2 KO than WT granule cells (WT = 102.6 ± 19.3%, n=6 vs. IB2 KO = 21.1 ± 8.7%, n=8; p=0.032; Fig. 6B). A possible explanation of this effect could be that granule cell intrinsic excitability was already increased in IB2 KO granule cells (cf. Fig.1B), such that the level of IB2 KO granule cell excitability in control was similar to that in WT granule cells after potentiation (Fig. 6B).

**Figure 6.**
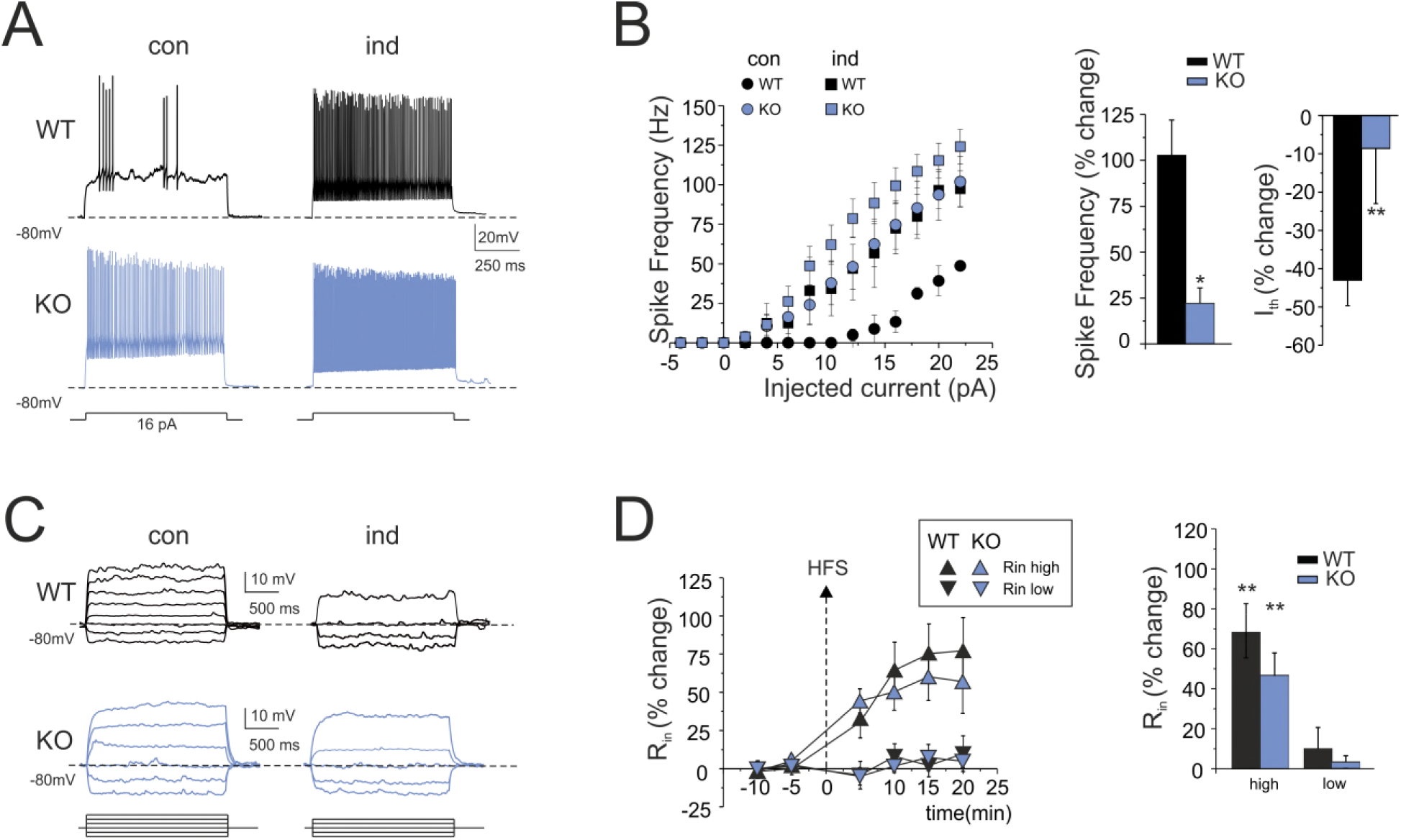
Long-term enhancement in granule cells intrinsic excitability. A. Voltage responses to current injection in WT and IB2 KO granule cells recorded in control and 20 min after HFS. Note that HFS enhances spike generation both in WT and IB2 KO granule cells.
B. Spike frequency is plotted as a function of current injection in control conditions and after HFS both in WT and IB2 KO mice. Note that, after HFS, spike frequency increases more in WT than in IB2 KO mice. The histograms compare the average spike frequency and threshold current (I_th_) changes in WT and IB2 KO mice. Data are reported as mean ± SEM; ^∗^p<0.05, ^∗∗^p<0.01.
C. subthreshold voltage responses to current injection in WT and IB2 KO granule cells recorded in control and 20 min after HFS. Note that the voltage-response in the high-potential region is enhanced both in WT and IB2 KO granule cells.
D. The plot shows the average time course of input resistance (R_in_) changes after HFS stimulation in two subthreshold membrane potential regions, < −70 mV (R_in-low_) and > −70 mV (R_in-high_). After HFS, in both WT and IB2 KO granule cells, R_in-high_ but not R_in-low_ increased. The histogram shows the average R_in_ changes for WT and IB2 KO mice. Data are reported as mean ± SEM; ^∗∗^p<0.01.

As a further control, we monitored the apparent granule cell input resistance (Fig. 6C,D) by measuring the response to small current steps (causing about 10 mV potential changes) either below −70 mV (R_in-low_) or above −70 mV (R_in-high_) (Armano et al., 2000). After HFS, R_in-hig_h rapidly increased in both WT and IB2 KO mice, following a similar time course and remained potentiated throughout the recordings (at least 20 min after HFS; average time courses are shown in Fig. 6D). At 20 min after HFS, R_in-hig_h increase was 67.8 ± 16.5% (n=8) (p=0.0014) in WT and 46.9 ± 9.0% (n=10) in IB2 KO mice (p=0.00012). This change was likely to contribute to the increased intrinsic excitability in both WT and IB2 KO. It should be noted that R_in-low_ remained unchanged in both WT and IB2 KO, providing an internal control for recording stability (Fig. 6C,D).

### Different mechanisms of LTP expression in IB2 KO granule cells

LTP expression was first assessed by analyzing changes in EPSC amplitude, variability (CV), and paired-pulse ratio (PPR) (Fig. 7A). The paired-pulse ratio (PPR) of EPSCs is generally considered to reflect changes in the probability of transmitter release in a pair of stimuli (Zucker and Regehr, 2002), while the coefficient of variation (CV) of EPSCs is a readout of presynaptic variability of quantal transmitter release upon repeated stimulation normalized by the mean (Malinow and Tsien, 1990; Manabe et al., 1993). In the recordings used for PPR and CV analysis, after HFS, the EPSCs showed a significant increase in WT (18.2 ± 3.4; n=8; p= 0.012) and IB2 KO mice (106.8 ± 51.8%; n=5; p= 0.05), while PPR (interstimulus interval 20 ms) showed a significant reduction in WT (-19.6 ± 9.3 %, n=8; p = 0.033) but not in IB2 KO (-6.7 ± 3.3 %, n=5; p = 0.1). Interestingly, CV significantly decreased in both WT and IB2 KO (WT = −28.3 ± 6.7, n=12; p=0.002; IB2 KO = −30.0 ± 8.0, n=9; p = 0.012). The CV decrease suggested that neurotransmitter release was increased not just in WT (Sola et al., 2004) but also in IB2 KO mice, although with some difference (see below).

**Figure 7.**
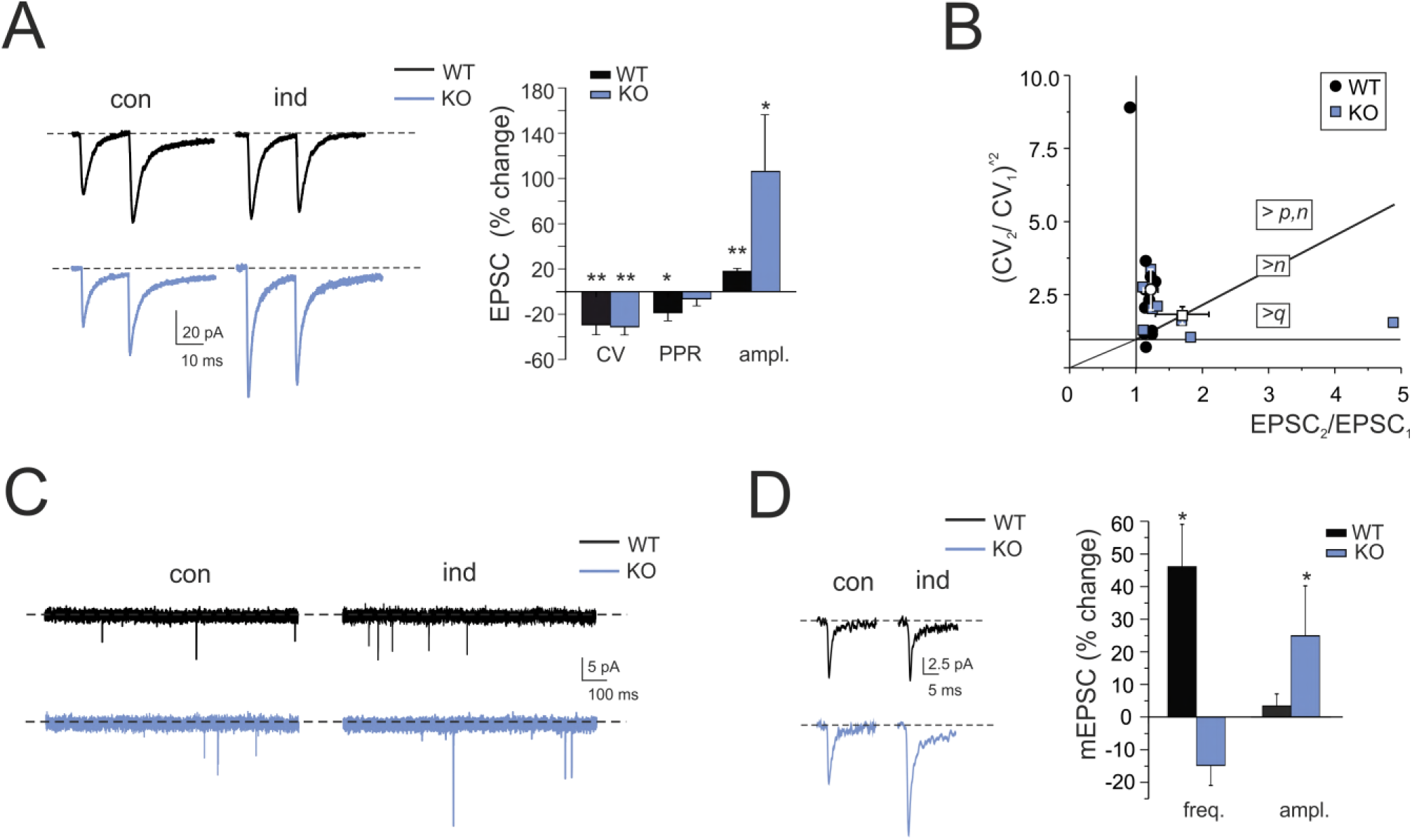
Mechanisms of LTP expression. A. EPSC_AMPA_ in WT and IB2 KO granule cells (average of 100 tracings in both cases) recorded in control and 20 min after HFS using paired-pulse stimulation (interstimulus interval 20 ms). The histogram shows the CV, PPR and EPSC_AMPA_ amplitude changes following HFS in WT and IB2 KO mice. Data are reported as mean ± SEM; ^∗^p<0.0, ^∗∗^p<0.01.
B. The (CV_2_/CV_1_)^−2^ *vs.* (EPSC_2_/EPSC_1_) plot shows that WT LTP points fall in the sector of increased quantal release (>*p*,*n*) while IB2 KO points fall on the diagonal (>*n*) and in the sector of increased quantum size (>*q*).
C. The traces show spontaneous synaptic activity before and after LTP induction in WT and IB2 KO granule cells. Following LTP induction, mEPSC frequency, but not amplitude, increased in WT while mEPSC amplitude, but not frequency, increased in IB2 KO mice.
D. Examples of individual mEPSCs before and after LTP induction in WT and IB2 KO granule cells. The histograms compare changes in mEPSC frequency and amplitude during LTP in WT and IB2 KO mice. Data are reported as mean ± SEM; ^∗^p<0.05.

The CV and PPR analysis cannot stand alone in determining the changes that could affect the neurotransmission process (Yang and Calakos, 2013). A further way to assess whether EPSC changes depend on the number of releasing sites (*n*), release probability (*p*) or quantum size (*q*) is to plot (CV_2_/CV_1_)^−2^ versus (M_2_/M_1_) (Bekkers and Stevens, 1990; Malinow and Tsien, 1990) (Fig. 7B). The WT experimental data points were distributed homogenously in the quadrant corresponding to *p/n* increase, with no point falling in the regions of a pure *n* or *q* change. Conversely, the IB2 KO experimental dataset was heterogeneously distributed over regions of *p*, *n* or *q* increase. These data distributions suggested that multiple presynaptic and postsynaptic mechanisms contributed to determine LTP at IB2 KO mossy fiber-granule cell synapses.

A second experimental approach to quantal analysis is to examine miniature postsynaptic currents (mEPSCs) before and after LTP induction (Fig. 7C,D) (Kullmann and Nicoll, 1992; Wyllie et al., 1994; Malgaroli et al., 1995). This method is especially useful at multi-quantal release synapses like here (Sola et al., 2004; Saviane and Silver, 2006) and can allow to distinguish between an increase in quantum content (*p* or *n*) or quantum size (*q*). Since here mEPSCs accounted for the whole spontaneous mossy fiber activity, in LTP experiments mEPSCs were recorded without TTX and were used to characterize the LTP expression mechanism (Sola et al., 2004). Moreover, in order to prevent mEPSC changes from being obscured by the contribution of non-potentiated synapses, we activated as many synapses as possible by raising stimulus intensity. Indeed, in these recordings, the EPSCs [(-59.0 ± 11.0 pA (n=4) in WT and −55.0 ± 14.9 pA (n=4) in IB2 KO mice] were about twice as large than those measured in Fig. 2 [(by comparison with single fiber EPSCs measured in similar recording conditions, this corresponded to activation of two to three mossy fibers (Sola et al., 2004)]. After HFS, the EPSCs increased (WT = 19.0 ± 2.0%, n=4; p = 0.02 vs. IB2 KO = 93.6 ± 49.7, n=4; p = 0.02) confirming larger LTP induction in IB2 KO than WT mice (cf. Fig. 7A). In the same recordings, mEPSCs amplitude did not vary in WT granule cells (3.3± 3.7 %, n=4; p = 0.4) but showed significant increase in IB2 KO granule cells (28.9 ± 5.66 %, n=4; p = 0.016). Conversely, mEPSC frequency showed a significant increase in WT granule cells (46.1± 12.9 %, n=4; p = 0.016) but did not show any significant changes in IB2 KO granule cells (- 16.9± 6.0 %, n=4; p = 0.11). Therefore, mEPSC analysis indicated that, while WT granule cells showed an increase in quantum content [(as reported previously in rats (Sola et al., 2004)], IB2 KO granule cells showed an increased quantum size.

In aggregate, these results confirm that LTP in wild type mice depends almost exclusively on increased neurotransmitter release probability (>*p*) and suggest that LTP in IB2 KO mice rests on a more complex mechanism including both changes in quantum content (>*p*, *n*) and quantum size (>*q*).

### Altered spatial distribution of LTP and LTD in the granular layer of IB2 KO mice

Given the enhanced LTP magnitude (cf. Fig. 5) and the altered C/S organization in IB2 KO granular layer (cf. Fig. 3), VSDi experiments were conducted in order to unravel possible alterations in the spatial distribution of LTP and LTD in IB2 KO granular layer. As recently shown using the same technique, the spatial distribution of areas undergoing LTP and LTD in the cerebellar granular layer displays a C/S-like organization, with LTP in the core and LTD in the surround (Gandolfi et al., 2015). The investigation of this feature in WT granular layer revealed a similar organization. Interestingly, the C/S organization of core-LTP and surround-LTD in IB2 KO granular layers showed a shape alteration with larger LTP cores and thinner LTD surrounds (Fig.8). The analysis of the granular layer areas with LTP and LTD revealed several abnormalities with respect to WT: i) LTP magnitude in the center was higher (WT = 28.4 ± 3.3% vs. IB2 KO =109.4±6.7%, n=6 for both, p=8×10^-6^); ii) LTP total area underwent an impressive increase (WT = 3.3 ± 1.5% vs. IB2 KO = 10.2 ± 3.3%, n=6 for both; p=0.047), iii) LTD total area and magnitude were decreased (WT = 91.4±1.9% vs. IB2 KO = 81.3±3.7%, n=6 for both; p=0.037; total LTD magnitude: WT = −34.9±2.8% vs. IB2 KO = −24.9±2.6%, n=6 for both, p=0.0026), and iv) the C/S shape showed a significant change in favor of LTP. In particular, the LTP-center was broader in IB2 KO compared to WT (core diameter: WT = 8.4 ± 0.7 μm vs. IB2 KO = 32.0 ± 4.1 μm, n=6 respectively; p=0.0005), and the LTD in the surround was less deep (WT = −37.3±1.5% vs. IB2 KO = −22.0±2.5%; n=6 for both; p=0.0004) (Fig. 8C,D).

**Figure 8.**
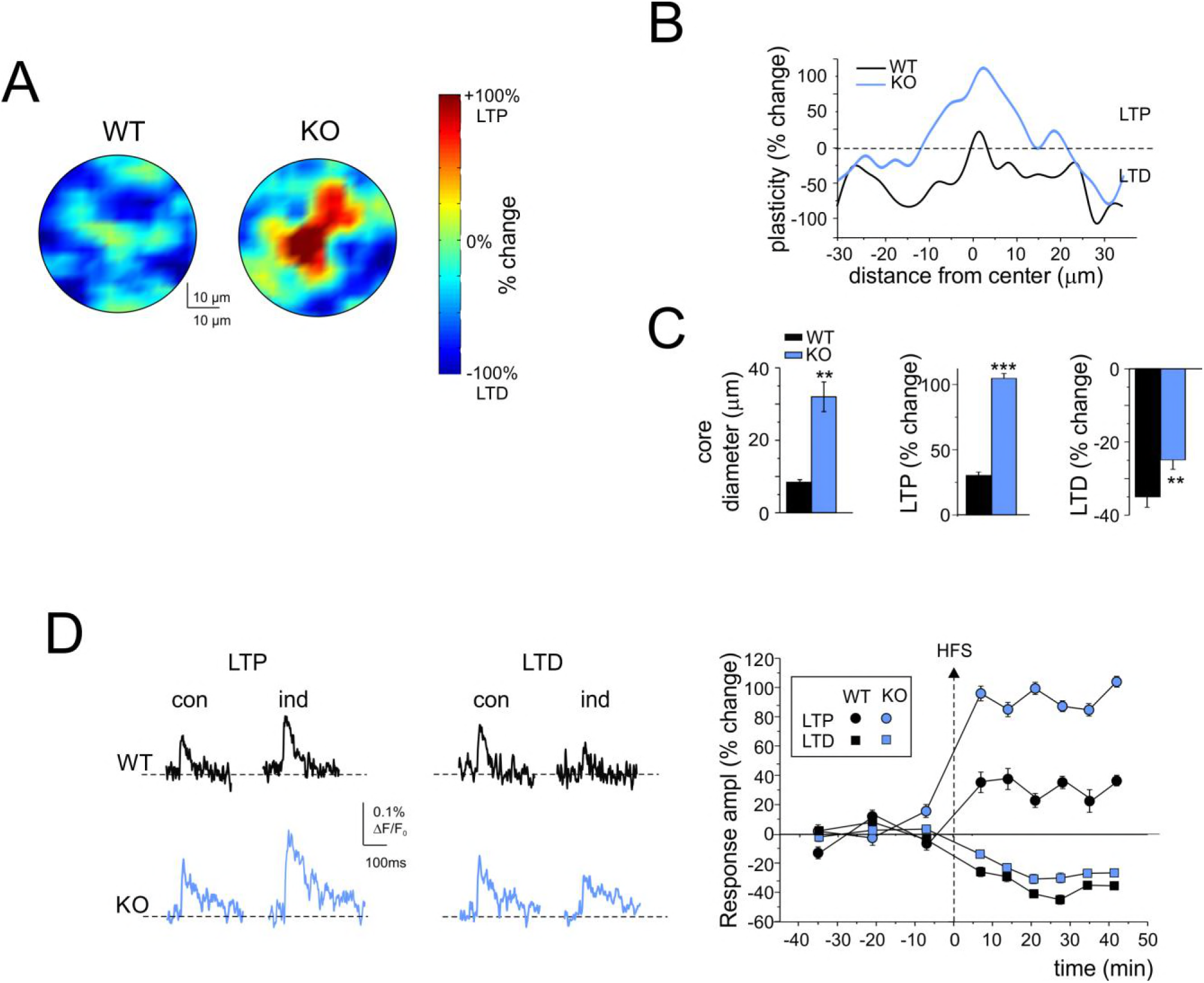
Spatial distribution of long-term plasticity of granular layer responses to mossy fiber stimulation. A. VSDi normalized maps showing the spatial distribution of LTP and LTD in WT and IB2 KO granular layers (average of 6 recordings in both cases).
B. The plot shows plasticity as a function of distance from the center for the maps shown in A. Note that in IB2 KO the LTP magnitude in the core is larger, and that the core is broader than in WT.
C. The histograms show, in WT and IB2 KO mice, the average core diameter and the LTP and LTD amplitude 30 minutes after HFS (n=6 for both). Note that the IB2 KO granular layer shows larger LTP smaller LTD and larger cores than WT. Data are reported as mean ± SEM; ^∗∗∗^p<0.001; ^∗∗^p<0.01.
D. VSDi recordings showing LTP and LTD of granular layer responses to mossy fiber stimulation. Exemplar traces before and 30 minutes after the induction protocol are reported for WT and IB2 KO. The plot shows the average time course of LTP and LTD for WT and IB2 KO (n=6 for both). Data are reported as mean ± SEM.

## Discussion

The main observation of this paper is that profound alterations in signal processing occur at the input stage of cerebellum in an ASD model, the IB2 KO mouse. Intrinsic excitability, synaptic transmission and synaptic plasticity in granule cells were enhanced in the absence of any compensation by the inhibitory circuit, causing a net increase in E/I balance. This in turn changed the spatial organization of neuronal responses, such that the core in C/S structures predominated over the inhibitory surround and LTP spread over larger areas.

### Granule cell hyper-functioning and the NMDA receptor-dependent current

In IB2 KO mice, cerebellar granule cells were hyper-functioning. *Enhanced synaptic transmission* appeared as a 2.4 times larger spike emission in response to high-frequency input bursts and was clearly correlated with larger NMDA receptor-mediated currents and increased intrinsic excitability. *Enhanced intrinsic excitability* appeared as a 2.1-7.1 (depending on current injection) higher efficiency in generating spikes during current injection and was correlated with larger Na^+^ and K^+^ membrane currents. *Enhanced synaptic plasticity* was manifest as a 5.3 times larger LTP compared to that normally measured at the mossy fiber - granule cell synapse (Prestori et al., 2008; Prestori et al., 2013). While normal LTP is almost entirely sustained by increased neurotransmitter release probability (Sola et al., 2004; D’Errico et al., 2009), IB2 KO LTP was expressed through a compound pre- and postsynaptic mechanism. This was consistently indicated by the increase in minis amplitude (>*q*) and decrease in EPSC PPR (>*n*, *p*) and confirmed by the ubiquitous distribution of points in the (CV_2_/CV_1_)^−2^ *vs.* (M_2_/M_1_) plot. The intervention of a postsynaptic expression mechanism was key to explain the neurotransmission increase in IB2 KO mice granule cells (~120%), which exceeds the theoretical limit of presynaptic expression alone (~60%; from (Sola et al., 2004)).

Interestingly, the whole set of alterations was likely to reflect, directly or indirectly, the NMDA receptor-mediated current enhancement occurring at the mossy fiber – granule cell synapse. In IB2 KO mice, the NMDA synaptic current of granule cells was increased by about 2.5 times, as anticipated by (Giza et al., 2010), while the AMPA receptor-mediated current was unaltered. During bursts, the granule cell NMDA current is known to exert a strong depolarizing action entraining a regenerative cycle (D’Angelo et al., 2005), in which depolarization removes NMDA channel unblock further increasing the NMDA current. The combination of this effect with enhanced intrinsic excitability could easily explain the enhanced synaptic transmission characterizing IB2 KO granule cells. In turn, enhanced NMDA receptor activation could also promote stronger plasticity of synaptic transmission and intrinsic excitability (Armano et al., 2000; Gall et al., 2005).

### Functional alterations of the granular layer microcircuit

Given the absence of changes in synaptic inhibition, the enhancements in excitatory synaptic transmission and intrinsic excitability provide an explanation for the remarkable increase in E/I balance, for the prevalence of core over surround in C/S responses and for the extension of the LTP territory. The C/S organization of the cerebellum granular layer depends on the balance between granule cell excitation and Golgi cell inhibition (Mapelli and D’Angelo, 2007). Here, the strong enhancement of the NMDA current could effectively counteract inhibition (Nieus et al., 2014) extending the core and changing the C/S from “Mexican hat” to “stovepipe hat” shape. The elevated input resistance and intrinsic excitability of IB2 KO granule cells could collaborate with elevated NMDA receptor-dependent transmission to spatially expand the excitatory footprint and zone of LTP. The consequences of NMDA receptor hyperfunctioning on the E/I balance and C/S changes could be further analyzed using realistic mathematical models of the granular layer (Solinas et al., 2010; Sudhakar et al., 2017).

NMDA receptor expression in granule cells is the strongest of cerebellum (Monaghan and Cotman, 1985) and is reasonable to speculate that a there damage could have a high impact on ASD pathogenesis. Although granular layer circuit alterations were uncompensated leading to a net E/I increase, some changes downstream might have a compensatory meaning. For example, in IB2 KO mice, the thinner molecular layer, the simplified dendritic tree and the smaller climbing fiber responses of Purkinje cells (Giza et al., 2010), may tend to limit the impact of granular layer overexcitation.

### Comparison of alterations with other circuits and ASD models

The alterations observed in the cerebellum granular layer of IB2 KO mice resemble in some respects those observed in other brain structures of ASD mice. An enhanced NMDA receptor-mediated neurotransmission was proposed to cause hyper-reactivity and hyper-plasticity in the somatosensory cortex (Rinaldi et al., 2007; Rinaldi et al., 2008b), in pyramidal neurons of the medium prefrontal cortex (Rinaldi et al., 2008c) and in the amygdala (Markram et al., 2008). Interestingly, hyper-reactivity and hyper-plasticity were correlated with enhanced E/I balance in relation with enhanced NMDA receptor-mediated neurotransmission (Markram et al., 2008). Therefore, our results support the concept that enhanced NMDA receptor-mediated neurotransmission is a common bottleneck for ASD pathogenesis in different brain areas, including the cerebellum. The change of C/S shape from “Mexican hat” to “stovepipe hat” is especially interesting in view of the ASD hypothesis developed for cortical minicolumns, the fundamental module of the neocortex (Casanova et al., 2002; Casanova et al., 2006; Hutsler and Casanova, 2016). The histological analysis postmortem of minicolumns in ASD patients has revealed reduced size and altered neuronal organization suggesting that lateral inhibition was reduced. In the C/S of the cerebellum granular layer, the *effectiveness* of lateral inhibition was indeed reduced by the increased intensity and extension of the excitation core. Therefore, a reduced effectiveness of surround inhibition of cortical and cerebellar modules may be a common trait of the disease in different brain microcircuits. The picture may be complicated by the interaction between causative, compensatory and developmental factors. For example, in *Gabrb3* mutants, an increased metabotropic glutamate receptor activation in deep cerebellar nuclei has been proposed to prevent the downstream propagation of effects and to protect from ASD in males (Mercer et al., 2016).

### Possible consequences of alterations on cerebellar functioning

The cerebellar granular layer has been proposed to perform *expansion recoding* and *spatial pattern separation* of input signals (Marr, 1969), which can be regulated by long-term synaptic plasticity at the mossy fiber - granule cell relay (Hansel et al., 2001; D’Angelo and De Zeeuw, 2009; D’Angelo, 2014). In IB2 KO mice, mossy fiber burst retransmission was enhanced and the effect could be further amplified by LTP (Nieus et al., 2006). Moreover, the excited areas were broader and poorly limited by surround inhibition. Therefore, expansion recoding and spatial pattern separation were likely to be compromised causing over-excitation of Purkinje cells and subsequent suppression of activity in deep-cerebellar nuclei. Altogether, these alterations could reverberate both on motor control (e.g. causing cerebellar motor symptoms) and on executive control (e.g. preventing novelty detection and attention switching) (D’Angelo and Casali, 2013), contributing to generate the combination of cerebellar and ASD symptoms presented by IB2 KO mice.

## Conclusions

The complex derangement of signal processing and plasticity in the cerebellum granular layer of IB2 KO mice supports a causative role of cerebellum in ASD pathogenesis. Microcircuit alterations resembled the *hallmarks* reported for cortical minicolumns, including synaptic hyper-reactivity, synaptic hyper-plasticity, increased E/I balance and C/S changes. In the cerebellum, these alterations have the potential of contributing to ASD as well as motor symptoms. Executive control may be affected by a dysfunction of loops connecting the cerebellum to associative (especially prefrontal) areas (Palesi et al., 2017), impairing novelty detection and attention switching and contributing to generate ASD symptoms (Schmahmann, 2004; Schmahmann et al., 2007; Schmahmann, 2010; D’Angelo and Casali, 2013). Future challenges will be to determine how cerebellar alterations combine and co-evolve with those occurring in other brain regions (Bolduc and Limperopoulos, 2009; Limperopoulos et al., 2009; Bolduc et al., 2011; Wang et al., 2014; Hampson and Blatt, 2015; Mosconi et al., 2015) and contribute to determine the different syndromic forms of ASD (Broussard, 2014; Hampson and Blatt, 2015; Mosconi et al., 2015; Zeidán-Chuliá et al., 2016).

## Acknowledgments.

This project has received funding from: the European Union’s Horizon 2020 Framework Programme for Research and Innovation under Grant Agreement No. 720270 (Human Brain Project SGA1); European Union grant Human Brain Project (HBP-604102); Fermi grant CNL to ED; Blue-Sky Research grant of the University of Pavia (BSR77992) to LM

